# Genetic entanglement enables ultra-stable biocontainment in the mammalian gut

**DOI:** 10.1101/2025.08.13.670093

**Authors:** Gary W. Foo, Aathavan S. Uruthirapathy, Claire Q. Zhang, Izabela Z. Batko, David E. Heinrichs, David R. Edgell

## Abstract

Imbalances in the mammalian gut are associated with acute and chronic conditions, and using engineered probiotic strains to deliver synthetic constructs to treat them is a promising strategy. However, high rates of mutational escape and genetic instability *in vivo* limit the effectiveness of biocontainment circuits needed for safe and effective use. Here, we describe STALEMATE (**S**equence en**TA**ng**LE**d **M**ulti l**A**yered gene**T**ic buff**E**ring), a dual-layered failsafe biocontainment strategy that entangles genetic sequences to create pseudo-essentiality and buffer against mutations. We entangled the colicin E9 immunity protein (Im9) with a thermoregulated meganuclease (TSM) by overlapping the reading frames. Mutations that disrupted this entanglement simultaneously inactivated both biocontainment layers, leading to cell death by the ColE9 nuclease and the elimination of escape mutants. By lengthening the entangled region, refining ColE9 expression, and optimizing the TSM sequence against *IS*911 insertion, we achieved escape rates below 10^*−*10^ as compared to rates of 10^*−*5^ with the non-entangled TSM. The STALEMATE system contained plasmids in *E. coli* Nissle 1917 for over a week in the mouse gastrointestinal tract with nearly undetectable escape rates upon excretion. STALEMATE offers a modular and simple biocontainment approach to buffer against mutational inactivation in the mammalian gut without a requirement for engineered bacteria or exogenous signaling ligands.

## Introduction

The composition and metabolic output of the microbiome of the human gastrointestinal tract are critical for health and development, and dysbioses or microbial imbalances are associated with acute and chronic health conditions^1, 2^. Traditional strategies to correct these imbalances are decreasing in efficacy, particularly for antibiotic-based therapies^3^. Engineered microorganisms and synthetic gene circuits offer a promising alternative to traditional approaches for human therapeutic use^4–6^, and also for applications in industrial, agricultural or environmental settings where microbial activities are critical^7–9^. Regardless of the setting, robust biocontainment strategies are of paramount concern for the safe and effective use of synthetic systems^10–20^. In particular, engineered microbes and synthetic systems designed for therapeutic use in the human gastrointestinal tract should ideally be regulated to restrict their growth to defined permissive conditions and to limit the spread of recombinant genetic material to native microbes. These concerns are particularly relevant for synthetic systems that use mobile genetic elements (MGEs), such as conjugative plasmids and transposons to propagate through microbiomes because they can circumvent chromosomal-based biocontainment systems by simply mobilizing to other microbes where they can persist for significant periods of time^21–26^.

Genetic instability and mutational escape are major issues when designing and implementing biocontainment systems, with current guidelines requiring an escape frequency of 10^−8^ or less^27^. Multi-layered containment systems based on auxotrophic dependencies or kill switches can exceed this guideline in laboratory-based conditions, but their reliance on external signaling molecules make implementation outside of laboratory conditions difficult^11, 12, 14, 17^. Installing multi-layered control systems on conjugative or other mobilizable delivery elements has shown promise in regulating spread, but high rates of mutational escape limit their efficacy. Deviations from optimal growth conditions are associated with elevated genetic instability and temperature fluctuations can upregulate insertion sequence (IS) transposition to increase *in vivo* rates of mutation^28–33^. More recently, synthetic sequence entanglements that link the expression of a biocontainment gene(s) with an essential gene to create a condition of pseudo-essentiality has shown promise in enhancing stability^34–37^. Complete sequence entanglements are not possible for all types of biocontainment systems, as recoding the primary sequence of each gene without adversely impacting or attenuating function can be problematic38, ^39^. Regardless of the approach, many biocontainment strategies rely on engineered bacterial strains that are not suitable as human therapeutics, or rely on chemical ligands that can be substrates for microbial metabolism, or that are not appropriate for human use^40–48^. Collectively, these limitations highlight the need to create biocontainment solutions that can reliably function outside of laboratory conditions and that are better suited to contain mobile genetic payloads.

We previously described a genetic safeguard composed of thermoregulated meganucleases (TSMs) designed for intracellular degradation of synthetic plasmid DNA in response to a temperature reduction outside of the mammalian gastrointestinal tract^31^ (Fig. 1A). The TSMs are composed of a LAGLIDADG homing endonuclease interrupted by a temperature-sensitive self-splicing intein that is active below 18°C. Transcription of the TSM is constitutive, but a functional TSM is reconstituted post-translationally by intein splicing at or below the permissive 18°C. Leveraging temperature as a single-input trigger to activate biocontainment eliminates the need for exogenous signaling ligands. We achieved escape rates of 10^−5^-10^−6^ with this system, but noted that insertion of IS*911* into the meganuclease sequence was the primary mechanism for mutational escape^31^ (Fig.S1).

**Figure 1.**
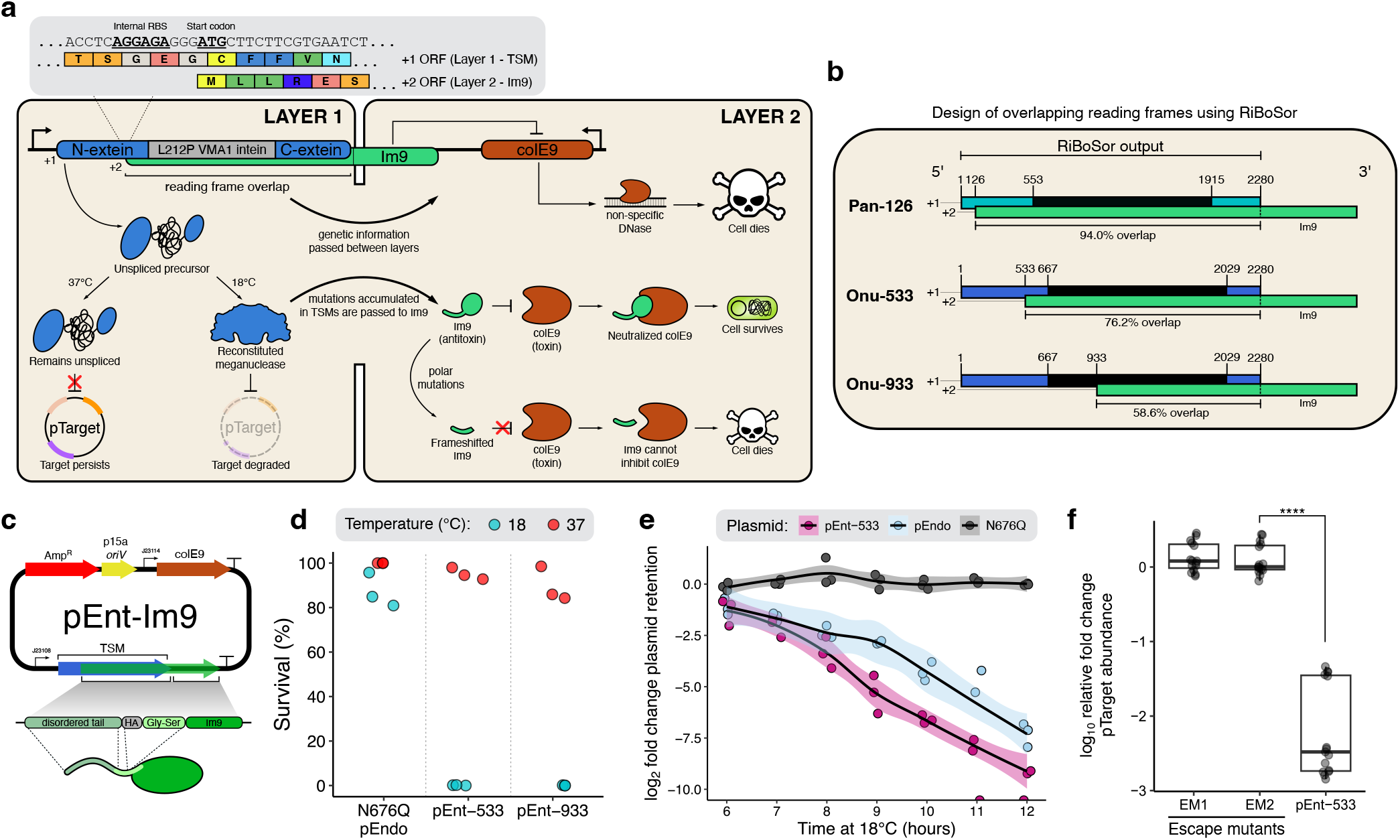
STALEMATE: Sequence entangled genetic buffers for biocontainment. **(A)** Schematic detailing the STALEMATE system. In the first layer, thermoregulated meganucleases (TSMs) degrade target plasmids when cells are incubated at 18°C. TSMs are genetically buffered in the second layer, via a sequence entanglement with the Im9 immunity protein. Polar mutations that occur in the +2 reading frame are usually lethal, by frameshifting Im9 and preventing the inhibition of the cytotoxic ColE9 DNase. Mutations that occur in the reading frame overlap affect both layers. **(B)** RiBoSor outputs using the the I-OnuI and I-PanMI TSMs as an input. Blue/cyan denotes extein sequences, and black denotes intein sequences with numbered start/end positions. Green denotes the +2 reading frame created by RiBoSor, with numbered start positions. **(C)** Plasmid map for pEnt-Im9. *oriV*, plasmid origin of replication; J23114/J23108, constitutive Anderson promoters; Amp^R^, ampicillin resistance gene; TSM, thermoregulated meganuclease; ColE9, colicin E9; Im9, ColE9 immunity protein. An HA-tag and Gly-Ser linked is included downstream of the RiBoSor tail to limit potential interference with Im9 folding. **(D)** Two-plasmid assay in *E. coli* Nissle 1917 using the entangled TSMs. Plasmid retention was determined as the ratio between colony forming units on LB agar with kanamycin compared to media without kanamycin. **(E)** Removal of pTarget by pEnt-533 compared to the unentangled I-OnuI TSM in *E. coli* Nissle. Each data point represents an individual biological replicate (n=3). **(F)** Intracellular degradation of pTarget confirmed by qPCR. EM1 and EM2 are two independently obtained escape mutants of pEnt-533. Data was obtained from three biological and five technical replicates, and each data point represents an individual replicate. Data are shown as box plots, with the bold line indicating the median, the rectangle the inter-quartile bounds, and the whiskers the data range. Statistical comparisons were performed with unpaired *t*-tests (\*\*\*\**P*<0.0001).

Here, we present STALEMATE (**S**equence En**TA**ng**LE**d **M**ulti L**A**yered Gene**T**ic Buff**E**ring) and use this strategy to create a failsafe system that dramatically improves the stability and significantly reduces mutational escape. STALEMATE uses a layered approach, entangling the primary biocontainment circuit (the TSM) with the colicin E9 immunity protein (Im9) by synthetic reading frame overlap to confer pseudo-essentiality and genetically buffer the TSMs from mutational escape (Fig.1A). Mutations that disrupt either open reading frame inactivate the protective function of Im9 and escape mutants are killed by the ColE9 nuclease. We found a 10,000-fold reduction in escape frequencies as compared to the original TSM. Plasmid-based STALEMATE systems showed enhanced stability and were maintained in the probiotic *Escherichia coli* Nissle 1917 strain without antibiotic selection for over 3 weeks. Altogether, the STALEMATE system and our data highlights a simple, modular, and effective approach to implementing pseudo-essentiality in a recombinant genetic circuit, without requiring bacterial genome engineering, chemical ligands to induce expression, or extensive primary sequence modifications to the entangled proteins.

## Results

### Design of the STALEMATE system

The plasmid-based STALEMATE system is composed of two layers with orthogonally functional biocontainment systems: a TSM in the first layer, and a sequence-entangled copy of the Im9 protein in the second (Fig.1A). The Im9 gene was cloned downstream of the TSM, and the Im9 +2 reading frame was extended upstream to overlap with different lengths of the TSM, depending on the construct. This +2 ORF extension created an N-terminal extension on the Im9 protein that was predicted to be unstructured by AlphaFold2 (Fig.S2). The ColE9 DNase gene was also cloned on the plasmid but in a different transcriptional orientation to the entangled TSM/Im9. While the TSM and Im9 are translationally independent, genetic information can be passed between the two layers in the overlapping region of the ORFs. Mutations in the overlapping region that inhibit TSM activity are passed onto the +2 Im9 reading frame to knockout Im9 function. We rationalized that Im9 inactivating mutations, or mutations that resulted in amino acid substitutions that impacted Im9 function, would no longer immunize cells against ColE9 nuclease activity. ColE9 would function as a failsafe mechanism to degrade all cellular DNA and cause cell death, eliminating escape mutants. Moreover, because the components of the STALEMATE system are transcribed constitutively at all temperatures, protection from mutational inactivation is not limited to the permissive temperature of 18°C for TSM activation.

This sequence entanglement setup differs from other entangled strategies as it limits the number of primary sequence changes needed to facilitate translation in each reading frame. We designed the first layer using the RiBoSor algorithm^35^ to codon optimize the I-OnuI and I-PanMI TSMs (Fig.1B, Table S3). A single missense mutation was required in the creation of all RiBoSor constructs in the +2 reading frame to create a ribosome binding site (RBS) or an AUG start codon. The internal ribosome binding site (AGGAGG) in pEnt-533 created a missense mutation that would have rendered I-OnuI catalytically inactive (E180G)^49–51^; this was avoided by using an alternative ribosome binding site (AGGAGA), which successfully restored enzyme activity while maintaining translation of the +2 ORF (Fig.S3). The other entanglements produced missense mutations that did not affect TSM protein activity.

### STALEMATE maintains activity in both reading frames

We confirmed that the activity of the +1 TSM ORF was not impacted by the introduction of the silent +2 ORF by cloning STALEMATE constructs onto a plasmid (pEnt) and performing two-plasmid cleavage assays with pEnt-533, pEnt-933, and a N676Q intein-splicing mutant (Fig.1C/D)^52^. Pan-126 was cytotoxic and not used in further experiments. Active TSMs should be functional below 18°C and cleave their cognate target site on pTarget, resulting in the loss of kanamycin-resistance (Fig.S4)^31, 53^. For these and all subsequent experiments, pEnt and pTarget were co-transformed into the probiotic *E. coli* Nissle 1917 strain, a generally recognized as safe (GRAS) bacteria for biotherapeutics in the mammalian gut^54^. We found robust TSM activity at the permissive (18°C) but not restrictive temperature (37°C) indicating that TSM thermoregulation and activity was maintained in the entangled TSM/Im9 construct (Fig.1D). As expected, no activity was found for the N676Q intein-dead negative control, consistent with previous data (Fig.1D)^31^.

Interestingly, we found that the STALEMATE-based systems cleaved pTarget more robustly than their unentangled counterparts. We performed *in vivo* time-point assays and found that the rate of plasmid clearance by pEnt-533 was faster than that observed for the unentangled pEndo (Fig.1E). We attribute this increased activity to be the result of increased stability of the STALEMATE system. Finally, we confirmed that the kanamycin-sensitive phenotype resulted from intracellular pTarget degradation by using quantitative PCR to measure plasmid levels. Active TSMs resulted in a 1000-fold relative reduction in the abundance of pTarget compared to two previously identified escape mutants of pEndo which were incapable of pTarget cleavage (Fig.1F, Fig.S5).

We next assessed gene expression in the +2 ORF using a chloramphenicol resistance gene. The chloramphenicol acetyltransferase gene was cloned downstream of the pEnt-533 TSM with a +2 ORF overlap of 1738 nts (580 residues) to generate pEnt-533-Cm^R^ (Fig.2A). When cells carrying pEnt-533-Cm^R^ were plated on solid media, growth was observed at 5 *µ*g/mL chloramphenicol but not at higher concentrations as compared to pEndo with an un-entangled Cm^R^ gene. This reduced chloramphenicol resistance could be due to the N-terminal tail on chloramphenicol acetyltransferase caused by the +2 ORF extension, or by the weakly active Anderson promoter (J23108) driving expression of the entangled TSM/CmR as compared to the native *cat* promoter on pEndo (Fig.2B). However, in liquid media, growth was observed at 25 *µ*g/mL chloramphenicol, albeit with a slower growth rate compared to pEndo (Fig.2C).

**Figure 2.**
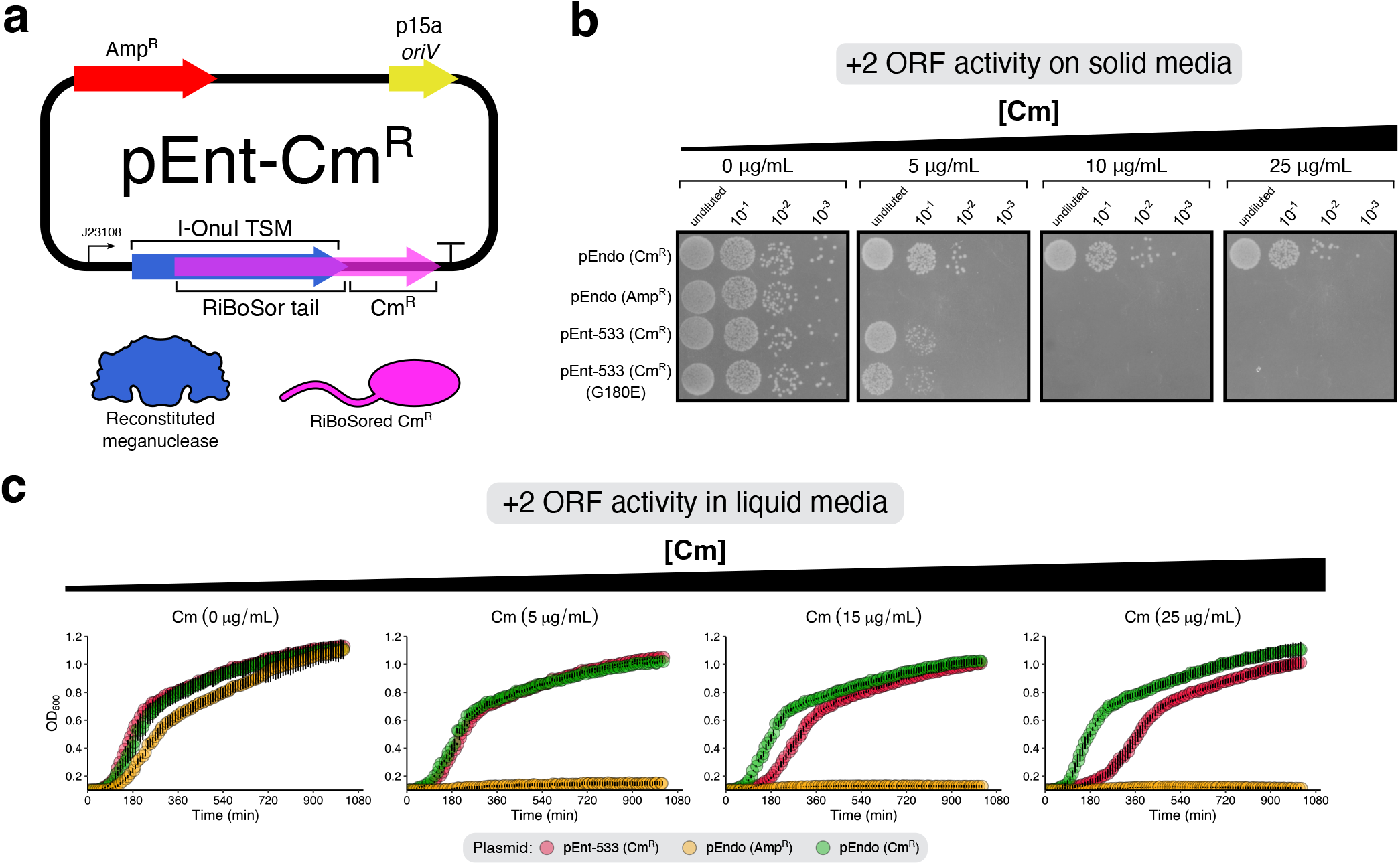
Chloramphenicol acetyltransferase activity in the second STALEMATE layer. **(A)** Plasmid map for pEnt-CmR. *oriV*, plasmid origin of replication; J23108, constitutive Anderson promoter; AmpR, ampicillin resistance gene; TSM, thermoregulated meganuclease; CmR, chloramphenicol acetyltransferase. The two products of the STALEMATE system on this plasmid are shown below. **(B)** STALEMATE-conferred resistance to chloramphenicol in *E. coli* spot plated on solid LB media containing different concentrations of chloramphenicol. pEndo (Cm^R^) is a version of the plasmid with the ampicillin resistance gene swapped out for chloramphenicol acetyltransferase without a RiBoSor tail, under the control of the native *cat* promoter. **(C)** Growth curves of *E. coli* carrying pEnt-Cm^R^ in liquid LB media containing different concentrations of chloramphenicol. Data points represents the mean and the whiskers represent the standard deviation (n=5).

Collectively, this data shows that the overlapping ORFs in the STALEMATE system are functional and that temperature-based regulation of the TSM first biocontaiment layer is not impacted by the sequence overlap. Moreover, the STALEMATE entangled TSM was more active than the non-entangled TSM, which we attribute to enhanced stability of the STALEMATE system.

### STALEMATE buffers against mutational inactivation

To create STALEMATE systems that do not rely on antibiotics, we swapped the chloramphenicol resistance gene with the Im9/ColE9-based second STALEMATE layer. Because ColE9 and Im9 are expressed at all temperatures, we anticipated that STALEMATE protection conferred by entangling TSMs and Im9 would not be limited to growth at 18°C. Mutations acquired during growth at 37°C or other temperatures would also be lethal and protect against escape mutants. Maintaining sufficient levels of Im9 to immunize against ColE9 is critical for functioning of the STALEMATE system and we first cloned ColE9 under the control of a weak ribosome binding site (BBa_B0031) and weak constitutive promoter (BBa_J23106) to attenuate ColE9 expression.

To test this setup, TSM activity was induced by overnight growth at 18°C and plated on media with and without kanamycin supplementation to determine the escape frequency. We observed escape frequencies of <10^−5.5^ and <10^−6^ with pEnt-533 and pEnt-933, an ~ 10-fold reduction in escape frequency as compared to the non-entangled pEndo (Fig.3B). To determine if the STALEMATE system genetically buffered the TSM from mutational inactivation, we sequenced 96 escape mutants of pEnt-533, pEnt-933, and pEndo. IS*911* transposition remained the primary mechanism for mutational escape, with 79.2%, 94.9%, and 94.9% of escapees resulting from IS*911* insertion for pEnt-533, pEnt-933, and pEndo, respectively (Fig.3C/D, Table S4). We ruled out chromosomal mutations as contributing to escape mutants by re-transforming plasmids into *E. coli* Nissle 1917 cells carrying pTarget (Fig.1F). In each case, escape mutant plasmids failed to contain pTarget, confirming plasmid loss-of-function rather than a chromosomal mutation. Moreover, Oxford Nanopore sequencing revealed that read lengths matched the expected sizes for plasmids, both with and without IS*911* insertion (Fig.S6).

**Figure 3.**
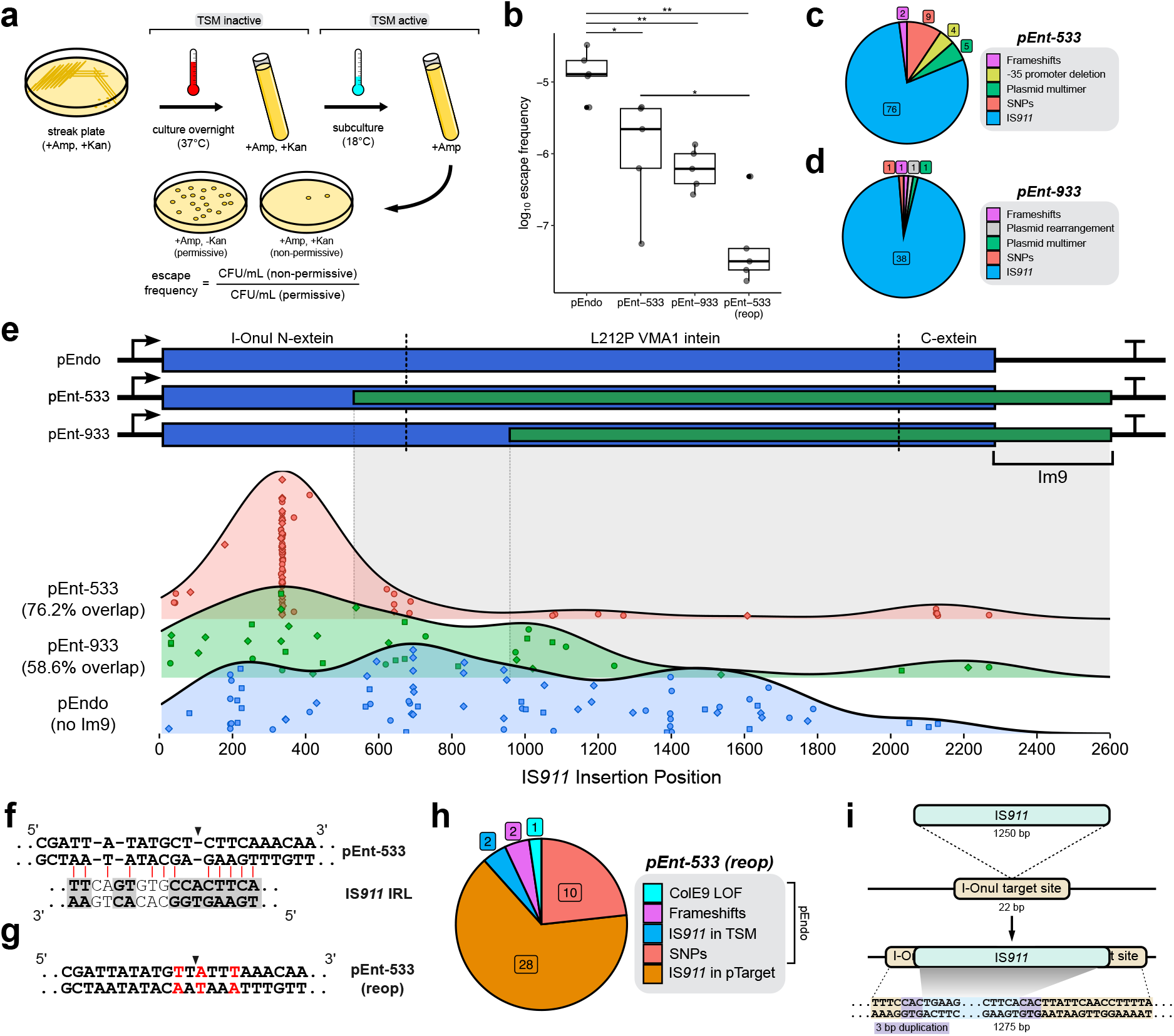
Alterations to the mutational landscape with Im9-based STALEMATE systems. **(A)** Schematic detailing how the escape frequency was determined. **(B)** Escape frequencies for pEnt-533 are improved over unentangled pEndo. Escape frequencies were determined as the ratio between escape mutants and total colony forming units. Each data point represents an individual replicate (n=5). **(C/D)** Pie charts categorizing the various methods of escape observed after full-plasmid sequencing of pEnt-533 (n=96) (C) and pEnt-933 (n=42) (D) escape mutants. **(E)** Ridge plots showing the distribution of IS*911* transposition events is biased towards the unentangled regions of the TSMs. Above is a schematic for the CDS of pEnt-533, pEnt-933, and pEndo, with dashed lines demarcating the intein/extein boundaries. Gray shaded areas show the extent of protection conferred by the entanglements. Each data point is an independently collected escape mutant (pEnt-533 (n=75), pEnt-933 (n=38), pEndo (n=75)) and shapes indicate the three different biological replicates from which the escape mutants were collected. **(F)** Sequence of the proposed IS*911* inverted repeat homology region in pEnt-533, aligned to the left inverted repeat of IS*911*. Black arrow shows the most commonly observed IS*911* insertion site in the pEnt-533 coding DNA sequence. Grey boxes show highly conserved regions in IS*911*’s left inverted repeat (IRL). **(G)** Sequence of the codon reoptimized version of pEnt-533. In red are the base pairs changed to reduce the preference for IS*911* transposition. **(H)** Categorization of escape mutants for the codon-reoptimized version of pEnt-533 (n=43). **(I)** Schematic detailing the new dominant mechanism for escape in pEnt-533 (reop). IS*911* now preferably transposes to interrupt the meganuclease cleavage site. Statistical analyses were performed with unpaired *t*-tests (\**P*<0.05, \*\**P*<0.01).

Notably, as the extent TSM/Im9 reading frame overlap increased, the distribution of IS*911* insertion sites dramatically shifted. This was most evident for the pEnt-533 construct, where 59/75 of IS*911* insertions mapped to the non-overlapped region of the TSM, as compared to the broader distribution of insertion sites in pEnt-933 or pEndo (Fig.3E). For pEnt-933 and pEndo, IS*911* insertions were distributed across the TSM DNA sequence and are consistent with the random or site-specific transposition mechanism of IS*911*^55, 56^. Among the analyzed escape mutants (253 in total), we did not find evidence for inactivation of ColE9, suggesting that excess Im9 limits leaky ColE9 activity^57^.

We also found 16 of 75 escape mutants with IS*911* transposition events in the TSM/Im9 reading frame overlap, all of which caused frameshifts in the TSM. Closer inspection of these events revealed that 14/16 resulted in a target site duplication of 3-bp and a net increase of 1253-bp that caused an Im9 frameshift (Fig.S7). The remaining events (2/16) resulted in a 4-bp duplication with a net increase of 1254-bp that maintained the Im9 reading frame (Fig.S7). It is unclear how these insertion events generated escape mutants, as both events should disrupt Im9 function and sensitize cells to ColE9 nuclease activity.

### Removing IS*911* insertion hotspots reduces mutational escape

We noted hotspots for IS*911* insertion in all three TSM constructs, particularly for pEnt-533 between nts 333-336 of the coding region. These events were observed across three biological replicates and thus were not derived from a single founder event. We examined the nucleotide sequence of the pEnt-533 TSM in this region, and found that many IS*911* insertions were localized 5-bp downstream of a region of high similarity to the left IS*911* terminal inverted repeat sequence (Fig.3F). Introduction of silent DNA substitutions to reoptimize this region (Fig.3G, reop) reduced the similarity to the IR sequence and further lowered the escape frequency to ~ 10^−7^ as compared to the non-optimized pEnt-533 (Fig.3B).Interestingly, sequencing of pEnt-533 (reop) escape mutants revealed a noticeable change in the pattern of a IS*911* insertion, with only 2/30 events localized to the TSM (Fig.3H, Table S4). The remaining 28 IS*911* insertions mapped to pTarget that contained the TSM target site; insertions in pTarget were not observed with previous escape mutants for any construct. Of the 28 IS*911* insertions in pTarget from different biological replicates, all of them were mapped to the I-OnuI target site for the TSM and had 3-bp target site duplications (Fig.3I). These insertions would presumably prevent cleavage of pTarget by the I-OnuI TSM to facilitate escape.

Taken together, these data show that the STALEMATE system genetically buffers the TSM from mutational inactivation. Ablating a IS*911*-like IR sequence further buffered the TSM from insertional activation, and shifted both the distribution and types of escape events. These changes effectively lowered the TSM escape rate 100-fold to <10^−7^ from the 10^−5^ escape rate observed for the non-entangled and non-optimized pEndo.

### Increasing ColE9 expression reduces escape frequencies

The STALEMATE system relies in part on the stoichiometry of Im9/ColE9 to immunize cells against ColE9 activity. However, it is possible that some mutations in the TSM/Im9 overlapped region that inactivate the TSM but do not completely ablate Im9 expression would escape ColE9 killing because there would still be an excess of Im9 relative to ColE9 (Fig.4A). We hypothesized that this mechanism of escape could be prevented by increasing expression of ColE9 to create an imbalance in Im9/ColE9 stoichiometry in cells with mutations that do not completely ablate Im9 expression, effectively raising the barrier for escape mutations (Fig.4B).

**Figure 4.**
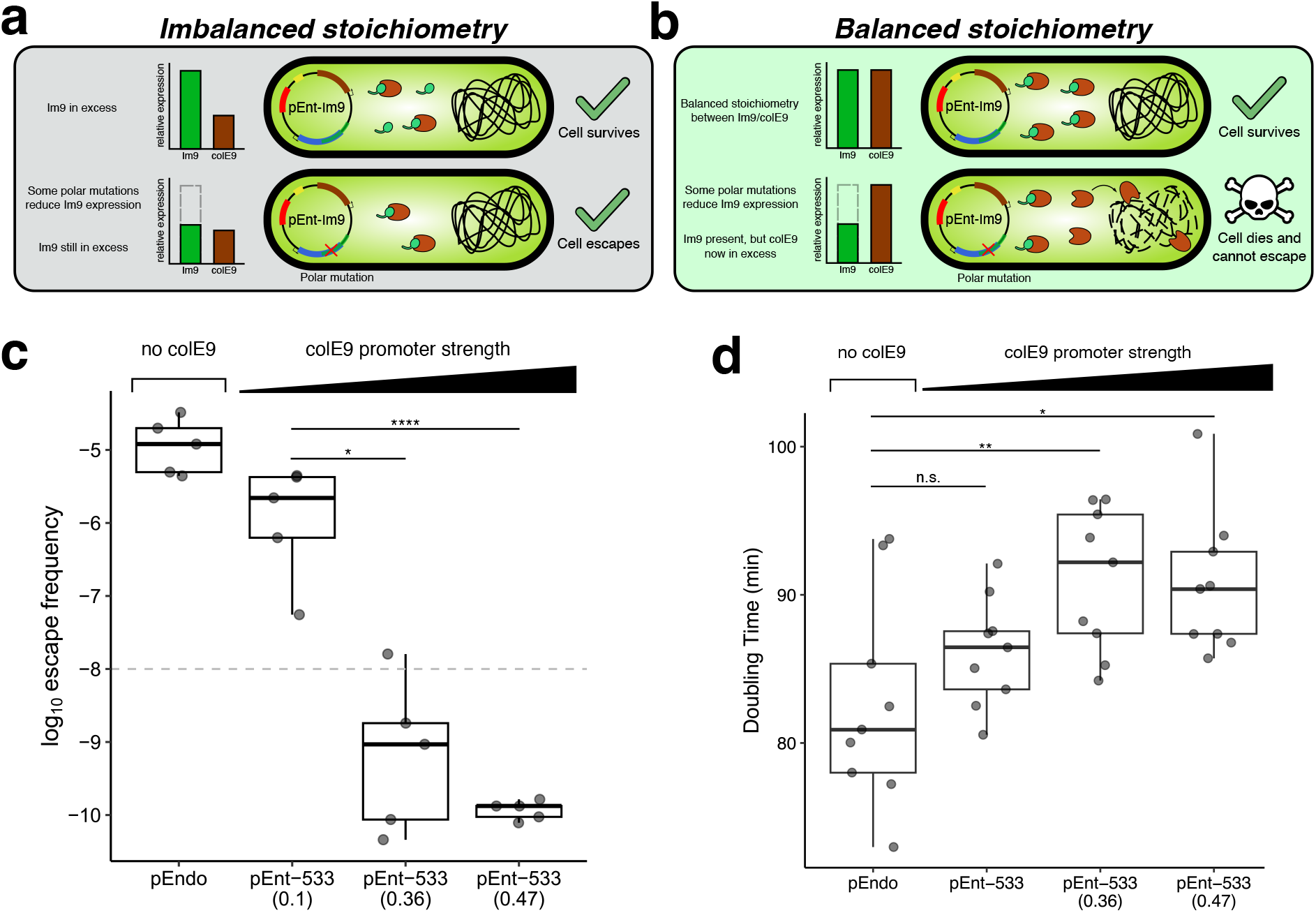
Effect of increasing ColE9 expression on escape frequencies. **(A)** Initial STALEMATE constructs possessed sub-optimal stoichiometries between Im9 and ColE9. Im9 is in substantial excess, therefore even minor perturbations to Im9 activity can be tolerated. **(B)** Increasing ColE9 expression closer to the maximum tolerable limit promotes more favourable stoichiometries between Im9/ColE9. Minor perturbations to Im9 are now more likely to be lethal. **(C)** Improved escape frequencies for pEndo and pEnt-533 using the stronger Anderson promoters driving ColE9 expression. The data is shown in box plots, and the escape frequency was determined by the ratio between the number of escape mutants and total colony forming units. Each data point is an individual replicate (n=5). Dashed line represents the minimum standard for biocontainment as per NIH guidelines. Statistical analyses were performed with unpaired *t*-tests (\**P*<0.05, \*\**P*<0.01, \*\*\*\**P*<0.0001). **(D)** Doubling times for pEnt-533 with stronger Anderson promoters driving ColE9 expression. Each data point is an individual replicate (n=9).

We tested this idea by increasing ColE9 expression relative to Im9 by using stronger Anderson promoter variants and measuring escape rates. We successfully cloned ColE9 under the control of the stronger J23107 (0.36) and J23106 (0.47) promoters, estimated to increase expression 4- and 5-fold relative to the J23114 promoter that was in our initial STALEMATE design (Table S5). pEnt-533 (0.36) and pEnt-533 (0.47) demonstrated significant improvements in escape frequency, with rates of 10^−9^ and 10^−10^, respectively (Fig.4C). However, we observed an upper limit for ColE9 expression, as a STALEMATE system using J23100 (1.0) that was cloned in *E. coli* EPI300 could not be transformed into *E. coli* Nissle 1917 and was not used for further experiments. We also observed that doubling times of *E. coli* Nissle with STALEMATE plasmids increased proportionally to ColE9 expression as compared to the original pEndo, suggesting that increased expression was a tradeoff for increased stability (Fig.4D).

Collectively, this data show that changing the relative expression of Im9 and ColE9 leads to significant improvements in the intrinsic stability of the STALEMATE system and reduces the escape rates 1000-to 10000-fold over non-entangled TSM. This level of escape exceeds the 10^−8^ NIH guideline by 100-fold^27^.

### STALEMATE systems are stable without antibiotic selection

Increasing ColE9 expression contributed to a 100-fold improvement in escape frequency but also raised concerns about long-term plasmid and strain stability. We rationalized that increasing ColE9 expression, while counterintuitive, may promote long-term plasmid stability without antibiotic selection by elimination of escape mutants where Im9 was inactivated at the single-clone level or at the population level by plasmid-policing (Fig.5A-C).^58–61^. To test this hypothesis, we passaged STALEMATE strains over three weeks by serial dilution, taking aliquots to measure both plasmid curing and escape rates (Fig.5D and E). As shown in Fig.5D, we observed that pEnt-533 (ColE9 0.47) maintained an escape frequency below the 10^−8^ NIH threshold for a two-week period. Strains carrying pEndo and pEnt-533 (ColE9 0.1) had escape frequencies in agreement with shorter-term measurements (Fig.5D and Fig.3B). Plasmid-curing data demonstrated that pEnt-533 had a similar rate to the vector-only control with a sharp decline in plasmid stability at day 18 (Fig.5E). In contrast, pEnt-ΔcolE9, which constitutively expresses the TSM/Im9 showed progressive loss over the 3-week period, with ~45% of cells losing the plasmid after three weeks. Collectively, this data indicates that a functional STALEMATE system does cause a significant selective disadvantage to cells and can be maintained in culture over ~ 3-weeks without antibiotic selection.

**Figure 5.**
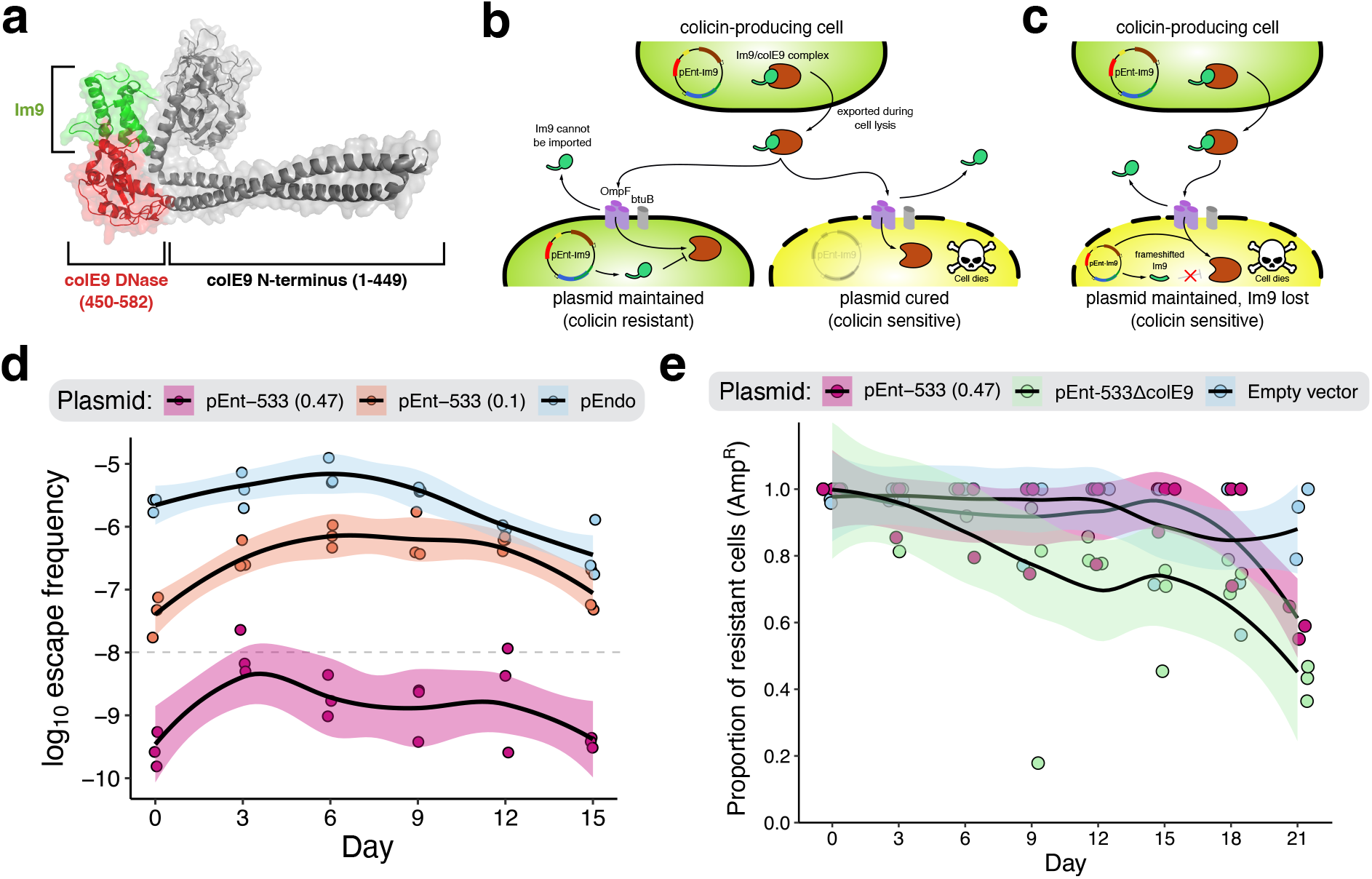
Colicinogenic entanglements promote plasmid stability at the population level. **(A)** Crystal structure of Im9 in complex with ColE9 (PDB:5EW5). **(B)** Schematic detailing plasmid policing at the population level, instigating death of cells in the population that spontaneously lose/cure the colicinogenic plasmid. **(C)** Schematic detailing policing against cells that maintain the plasmid, but lose/reduce expression of Im9 as a result of polar mutations in the reading frame overlap. Policing can occur either at the population level, or at the single-clone level. **(D)** Long-term escape frequency assay of pEnt-533 over 2 weeks demonstrates that stability persists. The escape frequency was determined by the ratio of escape mutants to total colony forming units. Each data point is an individual replicate (n=3), and the shaded areas represent 95% confidence intervals. Dashed line represents the minimum standard for biocontainment as per NIH guidelines. **(E)** Plasmid curing assays demonstrate reduced curing rate for pEnt-533 carrying ColE9 compared a version of the plasmid without ColE9. The proportion of resistant cells was determined by the ratio of colony forming units on selective LB media (100 *µ*g/mL carbenicillin) compared to non-selective LB. Each data point is an individual replicate (n=3).

### STALEMATE functions in the mouse gastrointestinal tract

We next determined whether the STALEMATE improvements to escape rates were maintained during passage through the mouse gastrointestinal (GI) tract, a commonly used model system for human GI-related disorders. In our experimental setup, pTarget should persist in the mouse gut because the 37°C temperature is a non-permissive temperature for TSM function in the first STALEMATE layer (Fig.6A). After elimination from the mouse gut and exposure to lower permissive temperatures, TSMs become active and cure pTarget. Because the TSM/Im9 and ColE9 are constitutively expressed at both 18°C and 37°C, this approach would allow detection of escape mutants that arose at any point during the experiment. We determined the escape frequency (and STALEMATE function) by plating bacteria from the feces of C57BL/6 mice at 18°C on kanamycin plates, and determined the total bacterial load by plating at 37°C (Fig.6B). We used the N676Q intein-splicing deficient TSM to determine a limit of detection for pTarget curing, finding ~ 10^6^ CFU/mg feces recovered on selective media each day of the experiment (Fig.6C). We also found similar total bacterial counts of ~ 10^6^ CFU/mg feces regardless of the pTarget or pEnt constructs, suggesting that the experiment did not adversely impact the mouse gut microbiome. Consistent with our previous data^31^, pEndo demonstrated a 4-log reduction in cells carrying pTarget at 18°C. This result confirmed TSM function in curing pTarget but also demonstrated the emergence of escape mutants. Crucially, *E. coli* Nissle carrying the STALEMATE plasmids pEnt-533 (0.36) or pEnt-533 (0.47) showed a 100-fold improvement over pEndo. We estimate the escape frequency of STALEMATE systems *in situ* to be between 10^−6^-10^−7^, a significant improvement over the 10^−4^ escape rate previously determined for pEndo^31^.

**Figure 6.**
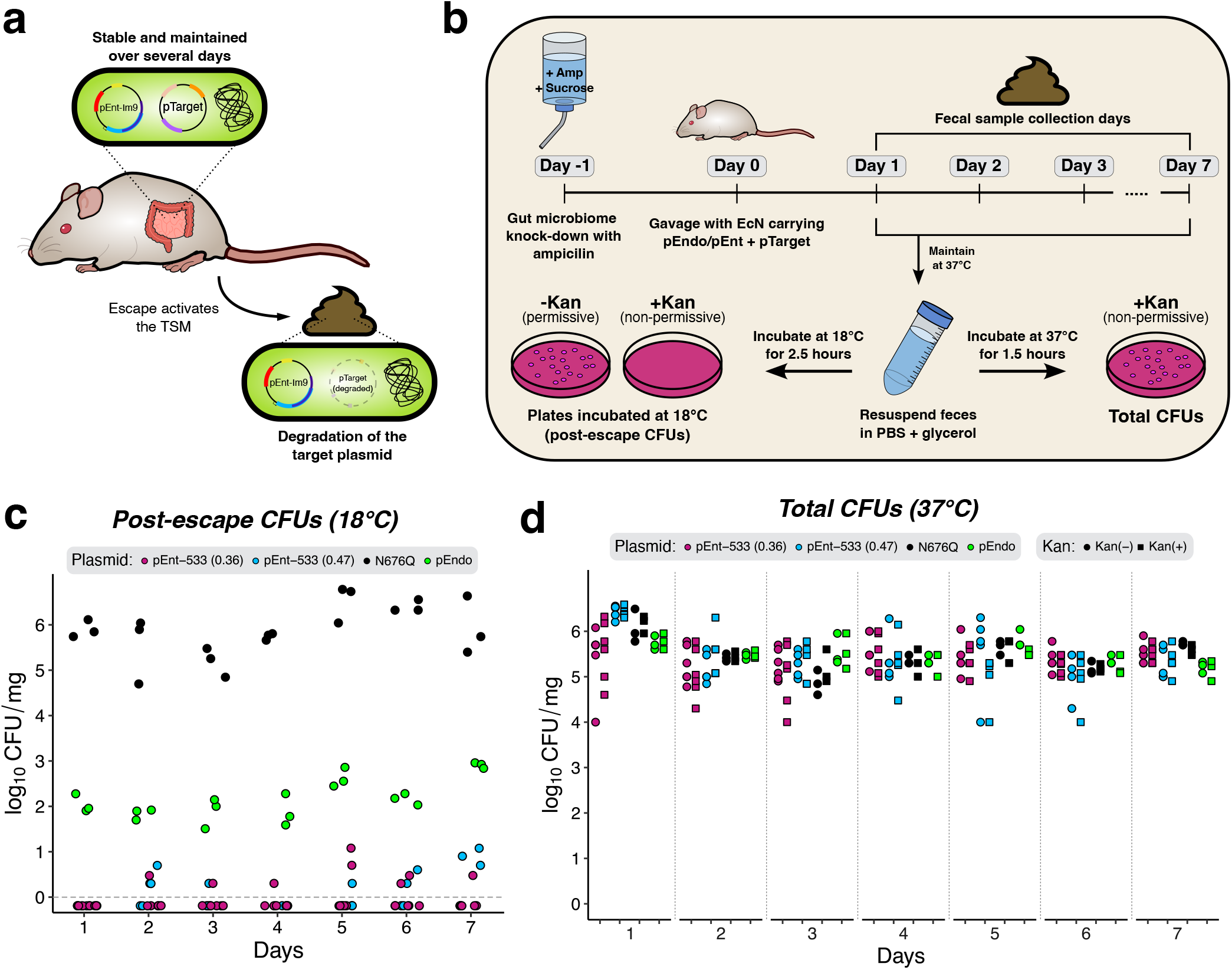
Improved stability of STALEMATE systems in the mouse gut over a week. **(A)** Schematic detailing the desired outcomes for pTarget retention, pre- and post-escape. **(B)** Schematic detailing the experimental setup for mouse model experiments. **(C)** Depletion of pTarget by pEndo or pEnt at 18°C. Colony forming units (CFU) from MacConkey agar supplemented with kanamycin were normalized to the mass of fecal samples. Dotted gray line denotes the limit of detection. **(D)** Maintenance of pTarget in the mouse gut at 37°C. Colony forming units (CFU) were obtained from fecal suspensions from MacConkey agar supplemented with-(Kan(+)) or without-kanamycin (Kan(−)) and were normalized to the mass of fecal samples. Each data point represents an individual replicate (pEndo (n=3), N676Q pEndo (n=3), pEnt-533(0.36) (n=5), pEnt-533(0.47) (n=6)).

## Discussion

In this study, we developed the STALEMATE system to promote pseudo-essentiality and enhance the stability of a biocontainment system installed on a mobilizable plasmid at both the cell and population level. Crucially, plasmids with STALEMATE systems can be maintained without antibiotic selection, a prerequisite for applications *in situ*. STALEMATE relies on synthetic sequence entanglements that are an emerging strategy towards implementing intrinsic genetic stability^35, 38, 62^. However, current entanglement approaches are difficult to implement for all proteins families, or proteins that are chimeric or bespoke in nature. In contrast, STALEMATE accommodates bespoke protein sequences with minimal primary sequence changes, preserving recombinant gene function by entangling sequences through open-reading frame extension^35^. The STALEMATE system developed here has two layered biocontainment modules, neither of which require engineering of the host genome and do not depend on exogenous chemical ligands to regulate gene expression. This genome-independent design ensures broad applicability across diverse organisms, and is particularly well-suited for the containment of mobile genetic elements that frequently transfer between microbes.

Kill-switches that rely on toxic genes are prone to mutational inactivation, requiring a delicate balance between evolutionary stability and robust biocontainment^63^. Even with tight repression, leaky gene expression often rapidly disables the safeguard, and kill-switch escape rates typically exceed those observed with auxotrophic containment systems^14, 17^. Our strategy of using the ColE9 bacteriocin in the second STALEMATE layer exploits the Im9 immunity protein to offset ColE9 cytotoxicity and enhance stability of the STALEMATE system. The system also acts to buffer the first biocontainment layer, the TSM from mutational inactivation and functions as a robust backup nuclease in the event of TSM/Im9 mutational escape; indeed, a single ColE9 molecule should be sufficient to kill a cell^59^. Our Im9/ColE9 system lacked the colicin lysin/releasing gene for the stress-induced release of ColE9 against competitor strains^61^. However, our data suggest that sufficient ColE9 is released to police Im9-lacking cells, possibly through spontaneous cell lysis during late stationary phase.

One aspect of escape that we explored in depth was mutational inactivation by endogenous insertion elements, specifically IS*911* where transposition is enhanced at low temperatures, including the permissive 18°C of the TSM biocontainment layer. However, the TSM/Im9 entanglements genetically buffered the TSM because IS*911* insertion or other types of mutations that occurred in the overlapping region would inactive the TSM and Im9, leading to ColE9-induced cell death. This setup allowed us to identify IS*911* insertional hotspots and to further buffer against insertion at these sites by re-optimizing the TSM/Im9 sequence. IS*911* is a member of the IS3 family of transposons that are widespread in Enterobacteriaceae and may be a source of mutational inactivation for synthetic constructs delivered to mammalian GI microbiomes^55^. Optimizing the DNA sequence of STALEMATE constructs to buffer against transposon inactivation could be incorporated into future STALEMATE designs based on the transposon composition of the target microbiome.

Interestingly, we did not find escape mutants with IS*911* insertions or other mutations that inactivated ColE9. This finding can be rationalized by the fact that ColE9 inactivation does not break the TSM biocontainment layer (unless a second mutation inactivated TSM/Im9) and clones with ColE9 knockouts would not be recovered on kanamycin plates. The plasmid-policing function of ColE9 would also restore an Im9/ColE9 system in a mixed population. However, we did find that an Im9+ColE9-strain experienced higher plasmid loss than a Im9+ColE9+ strain when grown in a monoculture. It is possible that plasmid loss is exacerbated by stress-inducing conditions in the mouse gut that would further complicate identification of ColE9 mutations.

Ideally, a complete sequence entanglement of TSM (in the +1 ORF) and Im9 (in the +2 ORF) would further buffer the system from inactivation. This may be possible with TSMs based on LAGLIDADG endonucleases other than I-OnuI or I-PanMI used here but is ultimately constrained by the tolerance of the +1 ORF to non-synonymous substitutions necessary to create an AUG start codon and ribosome-binding site for the +2 ORF^35^. Moreover, the length of the N-terminal extension created by a complete overlap could functionally impact the entangled ORFs. We found that the Pan-126 construct, with 94% of the TSM entangled with the +2 ORF extension, was cytotoxic at 18°C but that the cytotoxicity was not dependent on ColE9 function. While we do not understand how the sequence entanglement perturbs the I-PanMI TSM to create cytotoxicity, this shows that entanglements can lead to unanticipated changes in protein activity. For plasmid biocontainment purposes, we can also envision STALEMATE systems where the first biocontainment layer is based on a different site-specific endonuclease, possibly temperature-sensitive CRISPR variants or other site-specific nucleases. Moreover, for complete biocontainment of recombinant genetic material, we envision STALEMATE plasmids capable of their own removal, carrying the appropriate endonuclease cleavage sites to facilitate degradation of the plasmid in *cis* upon escape.

In summary, we developed the STALEMATE system to significantly reduce the mutational escape of biocontainment modules by creating a failsafe to kill escaping cells. In the context of plasmid biocontainment within the mammalian GI tract, our STALEMATE system does not affect the function of the primary biocontainment layer that targets plasmids for elimination at low temperatures outside of the GI tract. In laboratory conditions, our measured escape rates of <10^−10^ exceed the NIH guidelines 100-fold, and the escape rate estimated after passaging through the mouse GI tract of 10^−7^ is equal to or better than other strategies that rely on extensive genome engineering or multiple exogenous ligands for regulation. The STALEMATE system is compact, portable to different genetic contexts, and does not rely on exogenous signaling molecules for function.

## Methods

### Bacterial strains

*E. coli* EPI300 (F*′ λ*^*−*^ *mcrA* Δ(*mrr-hsdRMS-mcrBC*) *ϕ*80d*lacZδM15* Δ*(lac)X74 recA1 endA1 araD139* Δ(*ara, leu)7697 galU galK rpsL* (Str^*R*^) *nupG trfA dhfr*) (Epicenter) was used for plasmid cloning and storage purposes. *E. coli* Nissle 1917 (EcN) was used for all STALEMATE experiments.

### Plasmid construction

A list of primers is provided in Table S1 and plasmids in Table S2. Sequences of the Anderson promoters were from https://parts.igem.org/Promoters/Catalog/Anderson. All plasmids were assembled in *E. coli* EPI300 using either Gibson assembly or Golden Gate assembly^64, 65^. All Gibson assemblies were performed using the NEBuilder HiFi DNA Assembly kit (New England Biolabs, E2621), following manufacturer’s protocol. All Golden Gate reactions were performed with BsmBI-v2 (New England Biolabs, R0739) and T4 DNA ligase (New England Biolabs, M0202). Plasmids were designed in Benchling. Small oligonucleotides and large gene fragments (gBlocks) were ordered from Integrated DNA Technologies.

pEndo I-OnuI and pTarget are described in a previous study^31^. DNA sequences of the I-OnuI and I-PanMI TSMs used as an input to the RiBoSor algorithm are provided, in addition to the codon-reoptimized versions post-entanglement (Table S3). To make an empty vector, I-OnuI was removed from the original pEndo vector by inverse PCR, using DE-5792 and DE-7367. The resulting codon-reoptimized versions of pEnt-533, pEnt-933, and a copy of chloramphenicol acetyltransferase with a Gly-Ser tail were ordered as gBlocks with 30-bp homology overhangs, and cloned into the empty vector via Gibson assembly to produce pEnt-Cm^R^.

To swap out chloramphenicol acetyltransferase for Im9, Cm^R^ was removed from pEnt-Cm^R^ by inverse PCR, using DE-5792 and DE-7532. Im9 with a Gly-Ser tail was ordered as a gBlock with 30-bp homology overhangs, and cloned into the linearized vector by Gibson assembly to produce pEnt-ΔColE9 variant. To clone in ColE9, the pEnt vector was linearized again with DE-7362 and DE-7363, and a ColE9 cassette with the promoter (BBa_J23106), RBS (BBa_B0031), and double terminator (*rrnB T1*/*T7Te*) were ordered as a gBlock and cloned in via Gibson assembly. The promoter was swapped out by linearizing pEnt with DE-7363 and DE-8019, and oligonucleotides with the promoter flanked by 30-bp homology overhangs were cloned in via Gibson assembly. BBa_J23106 was cloned in with DE-8022, BBa_J23107 with DE-8021, and BBa_J23100 with DE-8026.

### Two-plasmid bacterial assays and escape assays

The two-plasmid bacterial assay was performed as previously described^31, 51, 66–69^. 50 ng of each pEndo/pEnt variant was transformed into 50 *µ*l EcN carrying pTarget. Cells recovered in 1 mL 2xYT media (16 g/L tryptone, 10 g/L yeast extract, 5 g/L NaCl) at 37°C and 225 rpm for 1 hour. Following the initial recovery, 1 mL of 2X induction media (2xYT, 200 *µ*g/mL carbenicillin) was added to the initial outgrowth. The media was then split, and half the outgrowth was induced at 37°C for 1.5 hours and the other half at 18°C for 2.5 hours to induce expression of the thermosensitive phenotype. After the induction period, media was serially diluted and spot plated or spread on 2xYT. Plates were incubated overnight at 37°C or 18°C for 5 days. Colonies were counted and plasmid retention was determined as the ratio of colonies grown on selective 2xYT media (100 *µ*g/mL carbenicillin, 50 *µ*g/mL kanamycin) compared to non-selective 2xYT media (100 *µ*g/mL carbenicillin). A splicing-deficient VMA1 intein mutant (N676Q) was used as a negative control.

Escape assays were performed similarly, with minor modifications. Rather than a transformation, EcN carrying either a variant of either pEndo or pEnt was grown overnight at 37°C under selection (100 *µ*g/mL carbenicillin, 50 *µ*g/mL kanamycin). The following day, the saturated culture was diluted 1:250 into 5 mL of non-selective LB (100 *µ*g/mL carbenicillin) and grown overnight at 18°C. On the third day, the culture was serially diluted and plated on selective- or non-selective LB media. The escape frequency was determined as the ratio of colony forming units on selective compared to non-selective LB plates.

### Time-point assays

EcN carrying either the I-OnuI TSM (pEndo), pEnt-533, or the N676Q intein splicing knockout were grown overnight at 37°C under selection. Overnight cultures were diluted 1:100 into LB media (100 *µ*g/mL carbenicillin), and incubation continued at 18°C for 12 hours. Aliquots of the culture were taken every hour, and plated on non-selective LB agar (100 *µ*g/mL carbenicillin) or selective LB (100 *µ*g/mL carbenicillin, 50 *µ*g/mL kanamycin). Plates were incubated at either 37°C or 18°C, and colonies were counted to determine the plasmid retention, calculated as previously described.

### Quantitative PCR

Protocol was performed as previously described^31^. After incubating for 12 hours at 18°C, 10 mL of culture was pelleted and resuspended in 500 *µ*L 1X phosphate-buffered saline (PBS). The resuspensions were boil-lysed at 95°C for 10 minutes, then immediately stored at −20°C. DNA concentration was determined with the Qubit 2 fluorometer (Life Technologies), and samples were diluted to 1 ng/*µ*L. Quantitative real-time PCR (qPCR) was performed using SYBR Select Master Mix (Applied Biosystems) on the Viia 7 Real-Time PCR system (ThermoFisher Scientific), amplifying a 150 bp region of the kanamycin resistance gene on pTarget using DE-7269 and DE-7270, and a 150 bp region of the *CspA* gene (ECOLIN_19660) on the chromosome of EcN using DE-7271 and DE-7272. Primer pairs were assessed for off-target activity by gel electrophoresis of the PCR reactions on a 1% agarose gel. Further validation was performed by melt curve analysis after running the samples. Three biological and five technical replicates were performed for each sample. Each reaction was performed in a total volume of 10 *µ*L, and included 1 ng of DNA and 400 nM of each primer. Thermocycler run parameters used the standard cycling mode: 50°C for 2 minutes, 95°C for 2 minutes, followed by 40 cycles at 95°C for 15 seconds and 60°C for 1 minute. Five replicates of a no-template control were also used for each of the primer pairs. Results were analyzed on the QuantStudio Software V1.3 (ThermoFisher Scientific). Data was plotted as the change in the quantity of pTarget relative to a catalytically inactive negative control. The relative quantity of pTarget was determined from standard curves produced by serial dilutions of purified pTarget and genomic DNA.

### Full-plasmid sequencing and IS*911* mapping

96 escape mutants were collected per sample, 32 escapees each from 3 biological replicates. All samples were collected from selective LB plates (100 *µ*g/mL carbenicillin, 50 *µ*g/mL kanamycin) grown at 18°C, colonies were picked and grown overnight in liquid selective LB media. Plasmids were extracted using the Monarch Plasmid Miniprep Kit (New England Biolabs), following the manufacturer’s protocol. Plasmids were barcoded using the Rapid Barcoding Kit 96 V14 (SQK-RBK114.96, Oxford Nanopore), following the manufacturer’s protocol. Samples were pooled and loaded onto a R10.4.1 MinION flow cell (FLO-MIN114, Oxford Nanopore), and left to run for 24 hours. The input library was assessed for sufficient concentration and quality with a Qubit 2 fluorometer (Life Technologies) and agarose gel electrophoresis. Reads were basecalled and demultiplexed using Dorado (v0.9.6). Reads were filtered by approximate size using fastcat (0.22.0), with a provided input size of 7500 bp. Reads were subsampled with Rasusa (v2.1.0) either to an expected coverage of 60x, or using the expected plasmid size to determine minimum coverage. Plasmids were assembled with Flye (2.9.5) and polished using Medaka (v2.0.1). Plasmids were assembled based on the expected sizes of pEnt (7.6 kb), pEndo (5.2 kb), and pTarget (3.7 kb). Samples that lacked sufficient read quality and depth to assemble plasmids were not used further. Assemblies were aligned to the pEnt-533 reference, and the locations of IS*911* was returned corresponding to the location on the TSM coding DNA sequence. If escape occurred through other means, they were also identified and recorded.

### Bacterial growth curves

EcN carrying pEndo, pEnt-533 (0.1), pEnt-533(0.36), and pEnt-533(0.47) were grown overnight at 37°C under selection. The following day, cultures were diluted 1:100 into 200 *µ*L LB media (100 *µ*g/mL carbenicillin) in a 96-well plate. Plates were incubated at 37°C, 225 rpm, double orbital shaking in the BioTek Epoch 2 Microplate Spectrophotometer, measuring the OD_600_ every 10 min for 18 hours. The doubling time was calculated via the formula: *doublingtime* = *ln*(2)*/ln*(1 + *growthrate*)

### Plasmid curing assays

EcN carrying an empty vector, pEnt-533(0.47), and a ΔColE9 variant were grown overnight at 37°C under selection (100 *µ*g/mL carbenicillin). The following day, cultures were passaged via a 1:100 dilution into fresh LB growth media without selection. This was repeated every day for 21 days. Every three days, an aliquot of the overnight culture was serially diluted and plated onto fresh selective LB (100 *µ*g/mL carbenicillin), or onto LB plates with glucose and without salt (10 g/L tryptone, 5 g/L yeast extract, 0.02% glucose) to prevent bacterial swarming. The proportion of resistant cells was the ratio between the CFU/mL on selective LB plates vs the anti-swarming plates.

### Mouse model experiments

Mouse model experiments were performed as previously described^31^. Three C57BL/6 female mice were kept per cage. Drinking water and feed were provided *ad libitum*. One day prior to gavage (Day −1), drinking water containing 2.5% sucrose was supplemented with ampicillin (1 g/L) to knockdown the gut microbiome. pEnt-533(0.36), pEnt-533(0.47), pEndo, and N676Q pEndo were previously transformed into EcN carrying pTarget and were grown overnight in selective LB at 37°C. On the day of gavage (Day 0), the overnight cultures were diluted 1:50 into fresh LB media, and grown to mid-log (OD_600_ ~ 0.5). The cells were pelleted and resuspended with 1X PBS, concentrating the cells to 10^8^ CFUs/100 *µ*L. Each mouse was gavaged with 100 *µ*L of the appropriate sample. For three days following gavage, mice fecal pellets were collected daily and resuspended in PBS (150 *µ*L/mg) by vortexing and mechanical agitation. Samples were serially diluted and plated on selective- (100 *µ*g/mL carbenicillin, 50 *µ*g/mL kanamycin) and non-selective (100 *µ*g/mL carbenicillin) MacConkey agar. Colonies were counted to determine the CFU/mg for each sample on the respective growth condition and temperature.

## Data Availability

BAM files from the Oxford Nanopore sequencing runs generated in this study were deposited in the Sequence Read Archive with the accession code PRJNA1293459. Other original data generated in this study is available as a supplementary data file. pEnt-533 (0.47) and pEnt-533-Cm^R^ have been deposited in Addgene as plasmid number 239788 and 239789, respectively.

## Supporting information

Mechanism and frequency of escape mutants in original TSM; AlphaFold2 predicted structures of STALEMATE N-terminal extensions; Sequences of entangled I-OnuI; Schematic of two-plasmid assay in *E. coli*; Standard curves for qPCR; Oxford Nanopore read length histograms; Alternative IS*911* target-site duplications; Tables of oligonucleotides and plasmids; Table of RiBoSor outputs Nfor entangled TSM/Im9 sequences; Summary table of escape mutants; Table of regulatory elements; GenBank formatted files of plasmids in this study.

## Author contributions

G.W.F., D.E.H., D.R.E. designed the experiments; G.W.F., A.S.U., C.Q.Z., I.Z.B. performed experiments; G.W.F., D.E.H., D.R.E. analyzed the results; G.W.F. and D.R.E. wrote the paper; D.R.E. and D.E.H. provided funding.

## Funding

This work was supported by a Project Grant (PJT 191939) from the Canadian Institutes of Health Research to D.R.E and D.E.H.

## Conflict of interest

The authors declare no competing financial interests.

## Acknowledgments

We thank members of the Edgell lab and Greg Gloor for reading the manuscript.

## Notes

### Competing Interest Statement

The authors have declared no competing interest.

