## Supplemental Data for "Genetic entanglement enables ultra-stable biocontainment in the mammalian gut"

### A temperature regulated mutationally-buffered biocontainment system for the mammalian gut

#### This document includes:

Figs. S1-S6

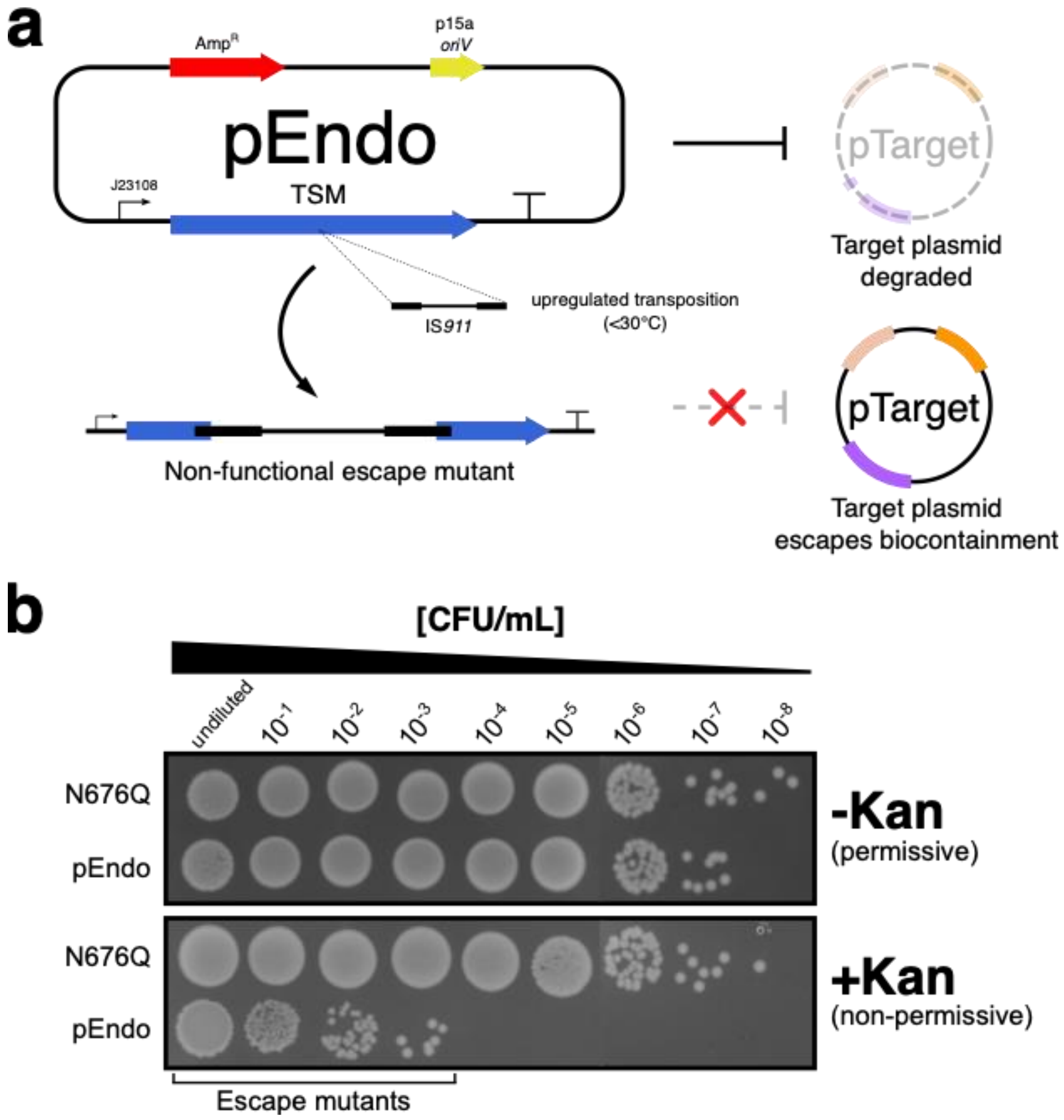


**Fig S1. Escape mechanism and frequency for the original I-OnuI TSM. (A)** Primary mechanism for circuit escape with the original TSMs. **(B)** I-OnuI TSM activity in *E. coli* on permissive and non-permissive LB plates, incubated at 18°C. Growth observed with the pEndo TSM on non-permissive media are considered escape mutants. The N676Q negative control is incapable of intein-splicing.

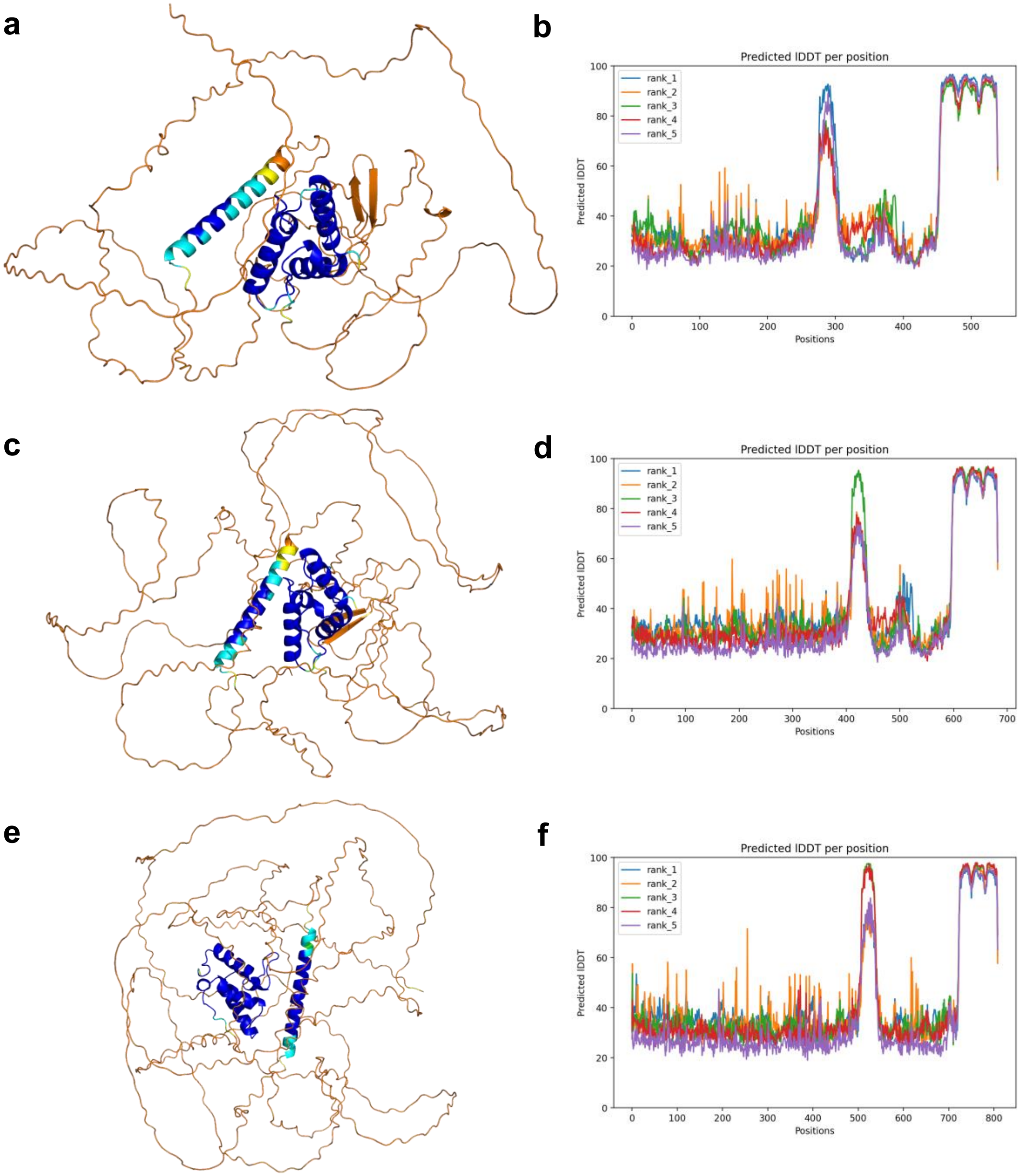


**Fig S2. AlphaFold2 predicted structures for STALEMATE constructs using lm9. (A,C,E)** Predicted structures for Onu-933 (A), Onu-533 (C), and Pan-126 (E) coloured by plDDT. **(B,D,F)** plDDT plots for Onu-933 (B), Onu-533 (D), and Pan-126 (F).

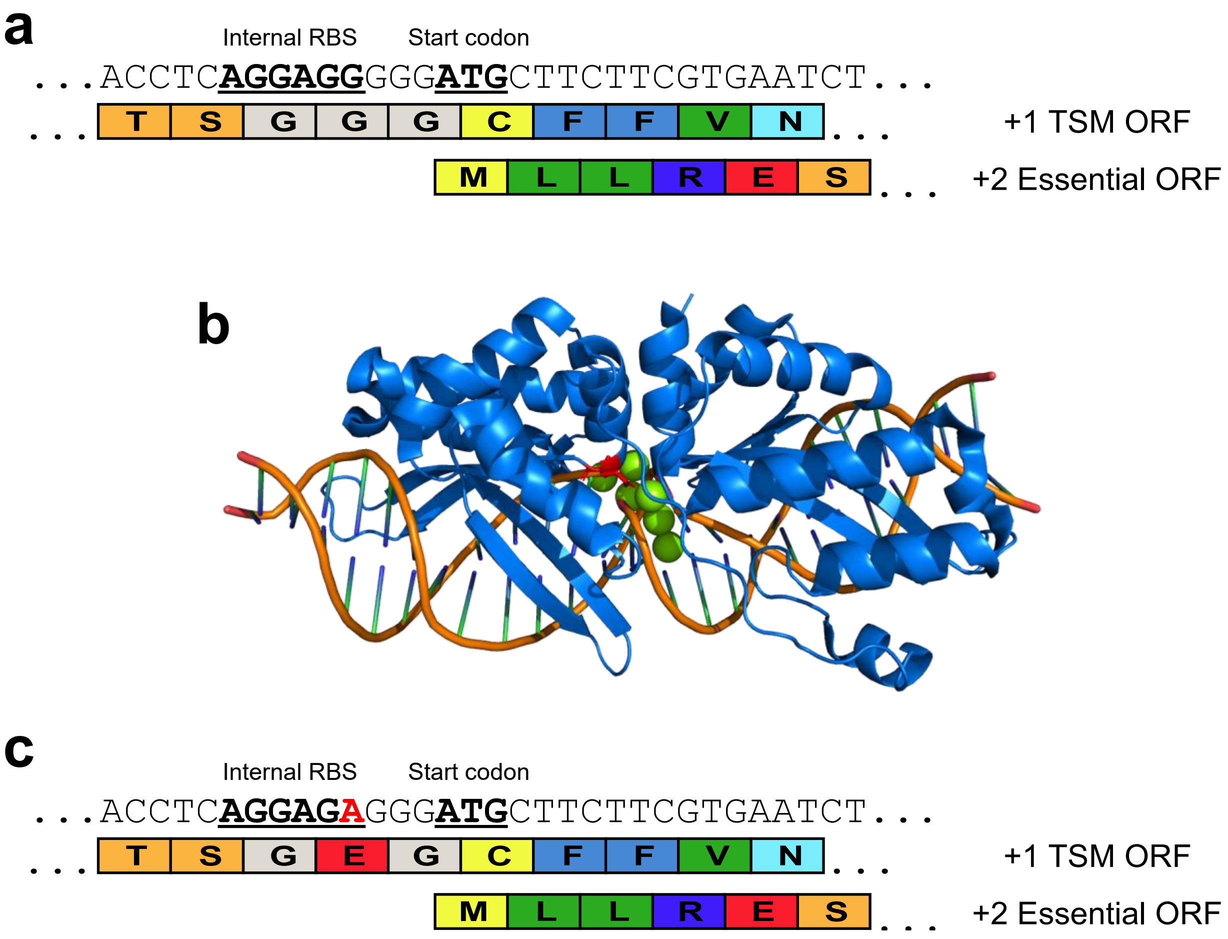


**Fig S3. Sequence of Onu-533 before and after restoring activity. (A)** The creation of the cognate AGGAGG ribosome binding site (RBS) in the +2 reading frame created the E180G catalytically inactive I-OnuI variant. **(B)** Crystal structure of I-OnuI (PDB:6BDA) in blue bound to its cognate target site. In red is the location of E180, one of the two residues coordinating the divalent metal ion cofactors shown in green. **(C)** Sequence of the corrected version of Onu-533, using an alternative RBS to restore E180.

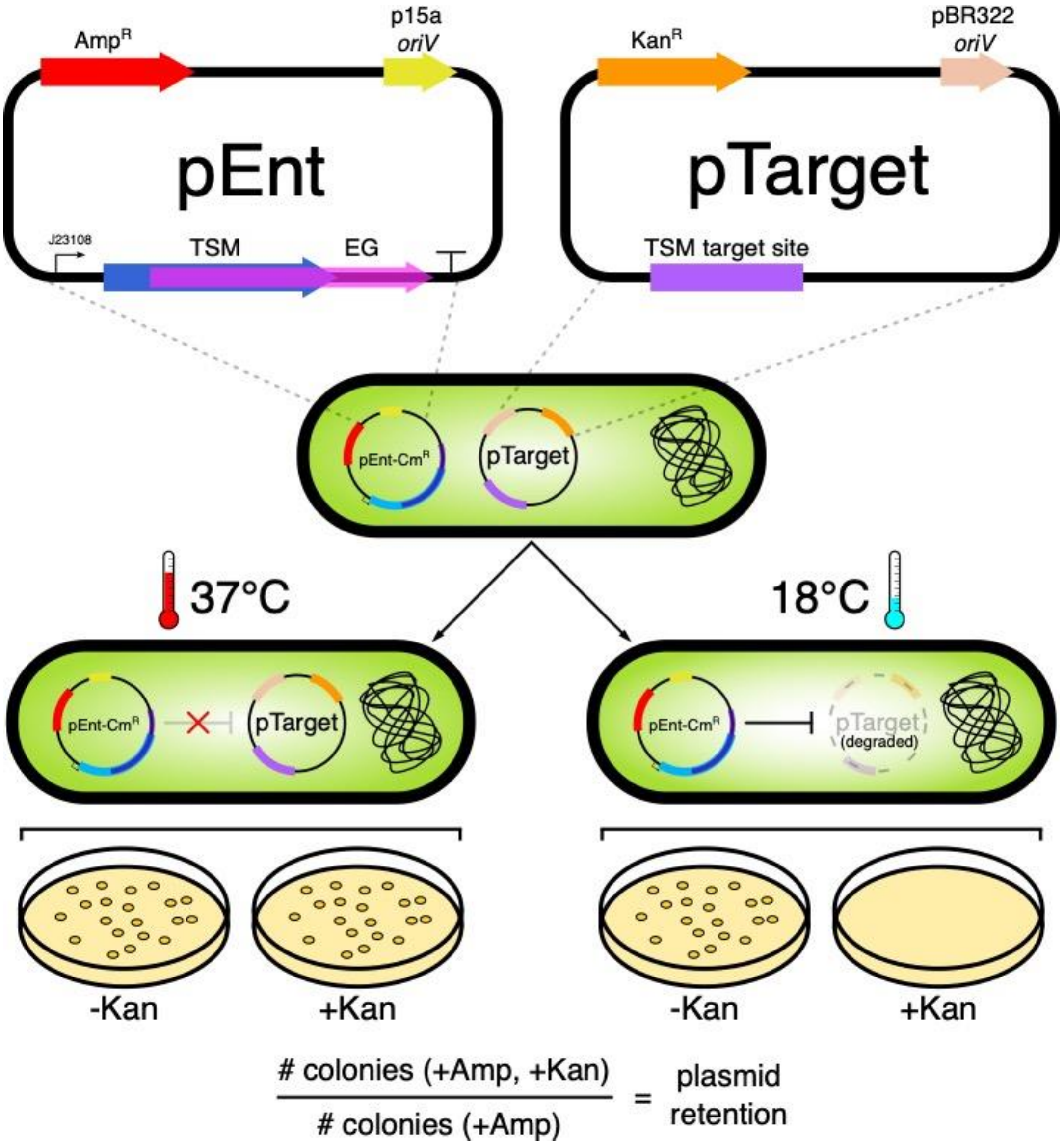


**Fig S4. Two-plasmid cleavage assay.** Plasmid maps for pEnt and pTarget used in this study. *oriV*, plasmid origin of replication; $Amp^R$, ampicillin resistance gene; $Kan^R$, kanamycin resistance gene; TSM, thermoregulated meganuclease; EG, essential gene; J23108, one of the Anderson promoters. The TSM target site is only cleaved when TSMs are reconstituted at 18°C and is measurable by the loss of kanamycin resistance on selective media.

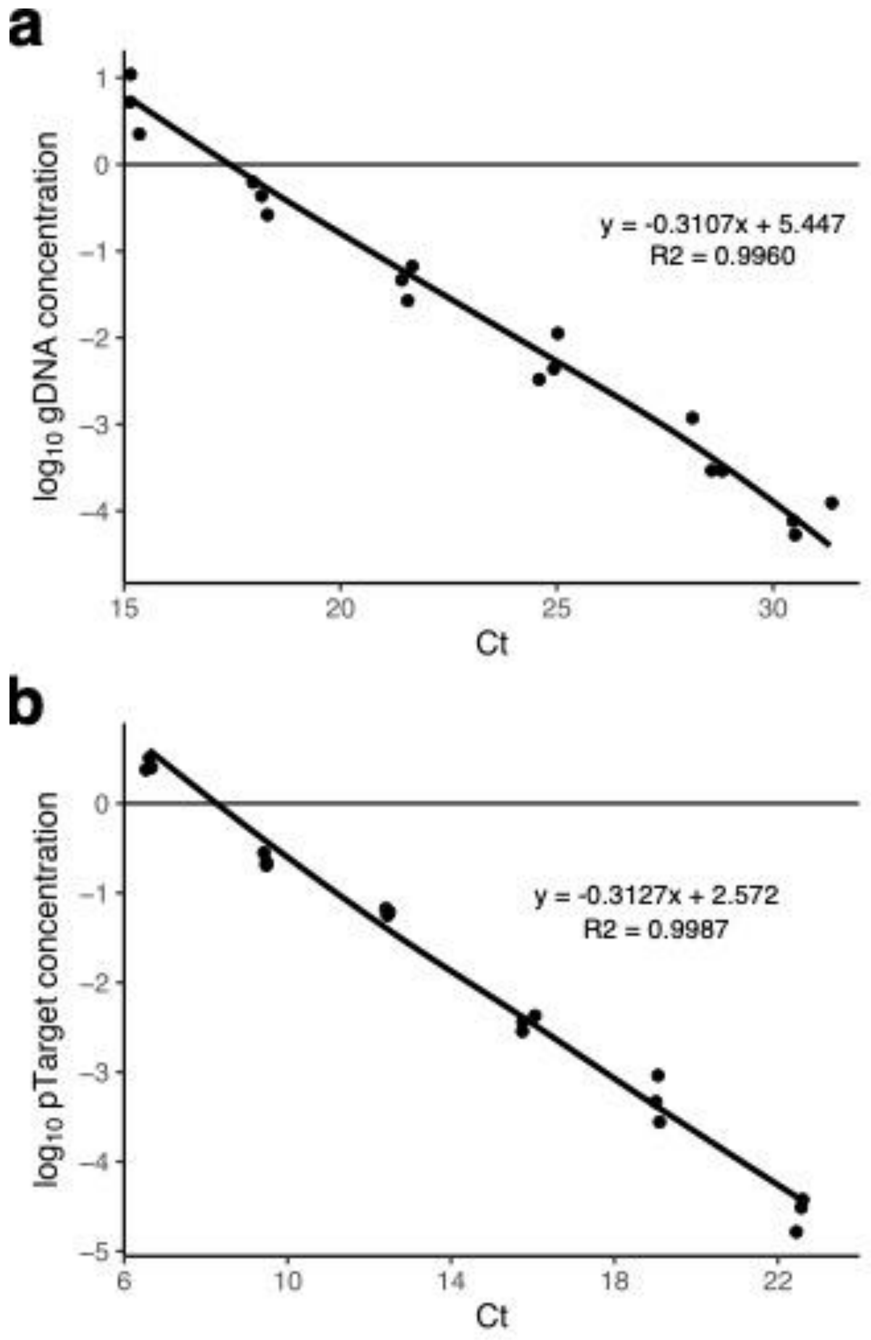


**Fig S5. qPCR standard curves for E. coli Nissle 1917 gDNA (A) and pTarget (B).** qPCR was performed with SYBR Select Master Mix (Applied Biosystems) on the Viia7 (ThermoFisher Scientific). The primer pair DE-7269 and DE-7270 were used against a 150 bp amplicon on the pTarget kanamycin resistance gene. The primer pair DE-7271 and DE-7272 targeted the *CspA* gene on the *E. coli* Nissle 1917 chromosome, also amplifying a 150 bp amplicon.

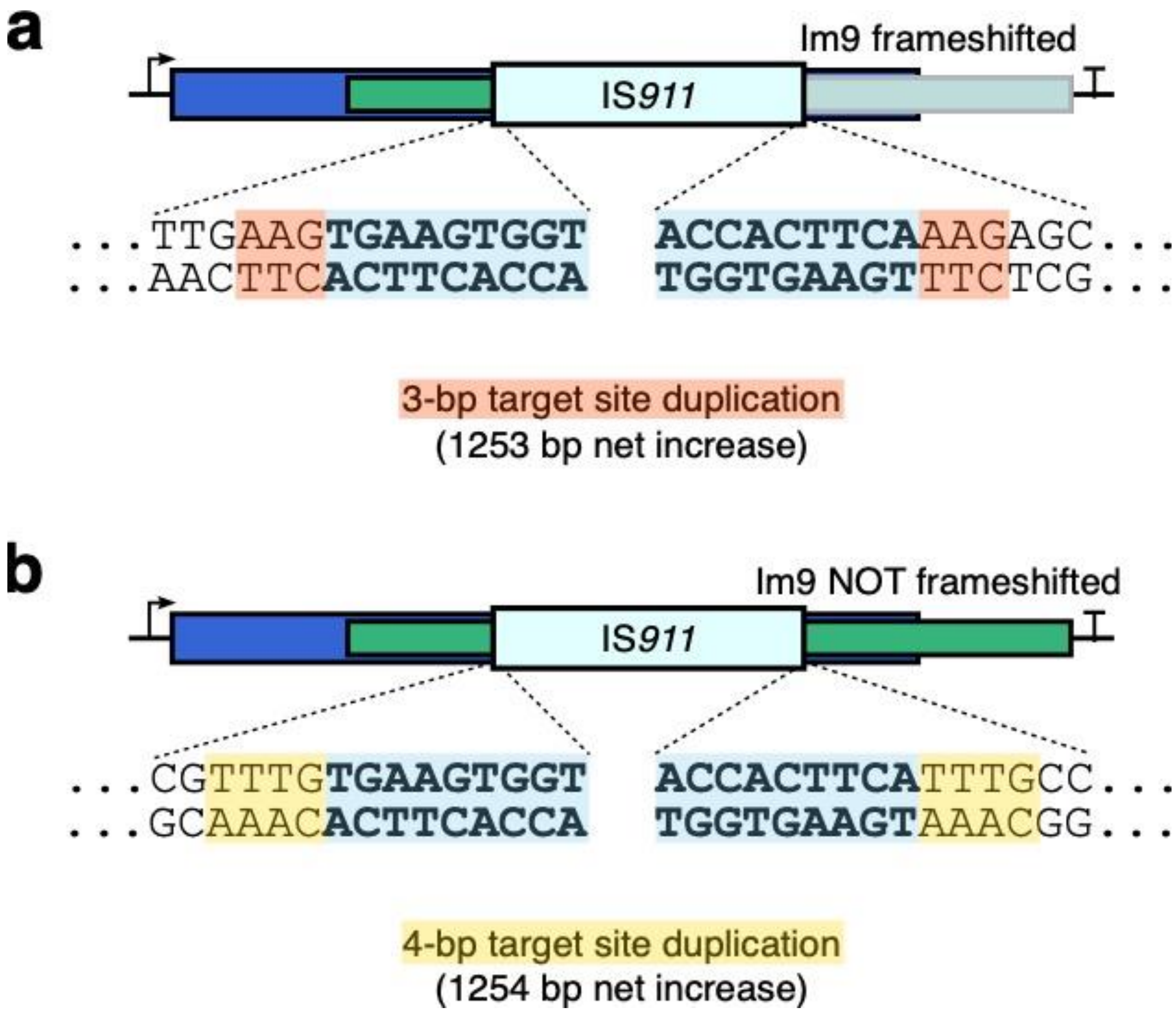


**Fig S6. Alternative IS*911* target site duplication as a mechanism for escape. (A)** Shown is the canonical 3-bp target site duplication that occurs during IS*911* transposition. IS*911* is 1250 bp long, and therefore results in a net 1252 bp increase, which causes a frameshift no matter the reading frame. **(B)** More rarely, a 4-bp target site duplication can occur and can result in a 1254 bp net increase. Depending on the orientation, this can result in a frameshift in one, but not the other reading frames.
